# Surface display of proximity labeling enzymes on extracellular vesicles for surfaceome and target cell mapping

**DOI:** 10.1101/2024.10.24.620083

**Authors:** Wenyi Zheng, Metoboroghene Mowoe, Wenqing Hou, Daniel W. Hagey, Koshi Imami, Samir EL Andaloussi

## Abstract

Extracellular vesicles (EV) surface proteins have important extracellular functions and determine cellular tropism; however, characterizing the EV surfaceome remains challenging with available methods. EV-mediated intercellular communication takes place primarily through interactions at the recipient cell membrane, underscoring the importance of methodological advances to map this interplay. Here, we leverage the proximity labeling enzyme APEX2 (Apurinic/Apyrimidinic Endodeoxyribonuclease 2) for high-fidelity analysis of the EV surfaceome and cellular tropism. Surface display of APEX2 on EVs is achieved through its genetic fusion with EV-sorting domains, such as CD63 and TSPAN2. Upon adding the substrates biotin-phenol and hydrogen peroxide, vesicle surface APEX2 enables biotinylation of EV integral and corona proteins as well as target cells in vitro. Further data mining of the EV surfaceome reveals potential scaffolds for the bioengineering of EVs. Altogether, we introduce a robust tool for EV surfaceome and target cell mapping and uncover novel EV-sorting domains for bioengineering.

## INTRODUCTION

Extracellular vesicles (EVs) are cell-derived particles with a plethora of physiological and pathological roles^1^, though the mechanisms by which they act are yet to be fully elucidated.^2,3^ The EV surface is composed of integral components from producer cells and a corona that becomes associated in the extracellular environment.^4^ Deciphering these molecular signatures is necessary to not only reveal their mechanisms of action, but also to identify effective diagnostic biomarkers.^5,6^ Furthermore, these surface proteins collectively guide EVs towards target cells.^4,7^ Besides their role in intracellular delivery,^8–12^ emerging evidence suggests that EVs may engage and signal target cells irrespective of cargo exchange.^13–21^ Thus, mapping the repertoire of EV-interacting cells could provide deeper insight into their biological and therapeutic roles. Achieving this requires the high-fidelity characterization of EV integral and corona components.

Despite the use of state-of-art purification methods, most commonly differential ultracentrifugation, EV preparations often contain considerable levels of non-EV proteins, such as albumin and lipoproteins.^22–24^ Moreover, the shear stress caused by sample centrifugation can lead to the dissociation and loss of EV corona proteins.^25^ Furthermore, available methods used to track EV tropism overwhelmingly rely on the cellular uptake of EVs^26^, and thus are unable to identify cells that respond to EVs through contact-dependent mechanisms. Therefore, an approach to identify EV integral and native corona proteins, as well as recipient cells, would be of broad utility.

Proximity labeling (PL) involves the use of engineered enzymes, such as peroxidases or biotin ligases, to generate a highly reactive, short-lived species that covalently tags neighboring molecules.^27^ Thus, PL enzymes have been tethered to EV surfaces to map the surfaceome and cellular interactome. They are either chemically conjugated to lipid anchors or lectins, for tethering with pre-isolated EVs,^28^ or genetically fused to the C1C2 domain that binds to phosphatidylserine lipids upon EV biogenesis^29^. However, these approaches are limited by PL enzyme dissociation from EVs^30^, the purity of EV preparations, catalytic activity levels^27^, and the vesicular display of phosphatidylserine^31^.

To date, APEX2 (Apurinic/Apyrimidinic Endodeoxyribonuclease 2) is the most active PL enzyme known.^27^ Using hydrogen peroxide (H_2_O_2_) as an electron donor, APEX2 converts the substrate biotin-phenol to biotin-phenoxyl radicals for the biotinylation of APEX2-proximal proteins. Subsequently, the biotinylated proteins may be isolated by streptavidin affinity purification and annotated using conventional mass spectrometry (MS) or other proteomic techniques. In this study, we genetically fuse APEX2 on the surface of EVs to map EV integral and native corona proteins, as well as target cells in vitro. Moreover, based on information from the EV integral surface proteome, we identify next-generation EV-sorting domains for bioengineering.

## RESULTS

### Selecting scaffold proteins for surface display of APEX2 on EVs

In previous studies, APEX2 has been terminally fused with domains-of-interest to highlight site-restricted expression; however, to the best of our knowledge, the vesicle surface display of APEX2 has yet to be reported. Here, we first designed engineering strategies for displaying APEX2 on the EV surface. The extracellular termini of single-pass transmembrane proteins (such as FPRP [Prostaglandin F2 receptor negative regulator], LAMP2 [lysosome-associated membrane glycoprotein 2], and PTTG1IP [pituitary tumor-transforming gene 1 protein-interacting protein]) have been shown to display cargo proteins^32–34^; thus, these strategies were tested in this study. In addition, since the N- and C- termini of APEX2 are in close proximityb^35^, we tested the integral fusion of APEX2 into the extracellular loop of tetraspanin proteins, CD63^36^ and TSPAN2^37^, assuming this would not influence protein folding and function. On APEX2, mNeonGreen (mNG) reporter protein was terminally fused to these constructs for intravesicular loading, to enable the detection of engineered cells and EVs. Wild-type CD63 without APEX2 was used as the negative control (Figure 2a). Thus, these six engineering strategies were compared for their efficiency in the surface display of APEX2.

**Figure 1.**
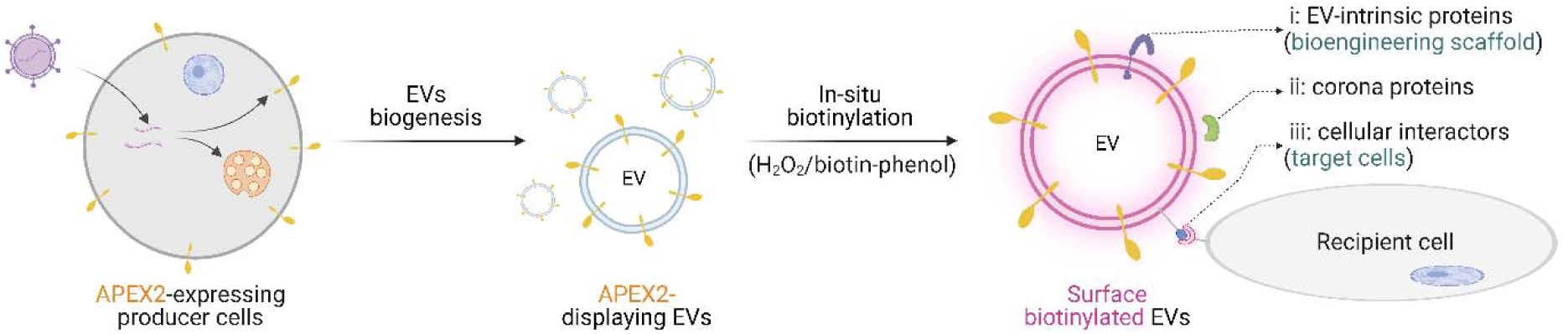
Proximity labeling using vesicle surface APEX2. EV producer cells are genetically engineered to express APEX2 on the EV surface. APEX2-catalyzed biotinylation of EVs enables the identification of EV integral and corona proteins of plasma-exposed EVs, as well as target cells. In addition, the EV integral surfaceome proves a valuable source of potential EV-sorting scaffold proteins.

**Figure 2.**
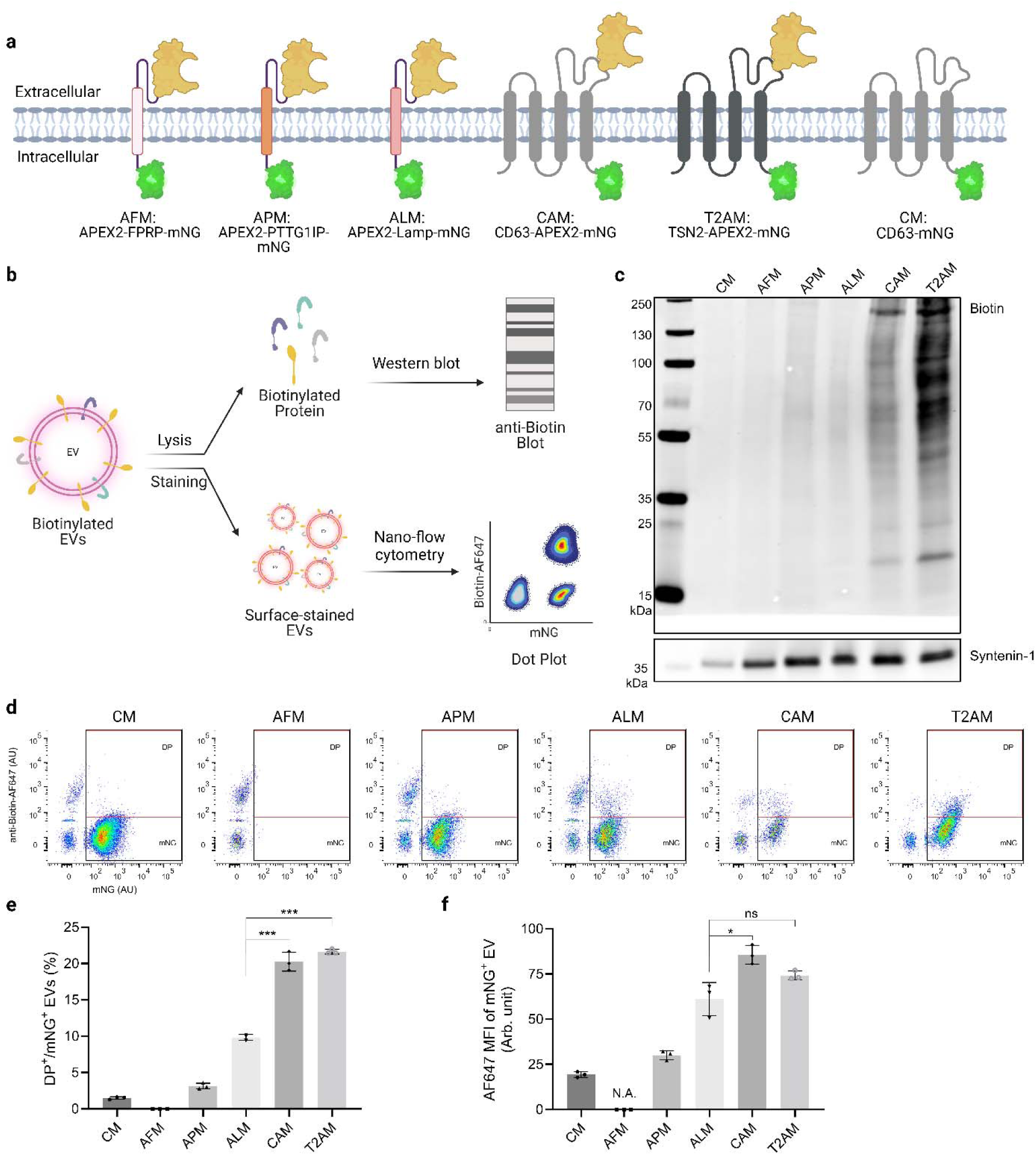
Comparing different scaffold proteins for surface display of APEX2. (a) Construct design for surface display of APEX2 on EVs. (2) Validation of vesicle surface APEX2 activity using western blotting and nano-flow cytometry. (c) Analysis of biotinylated proteins in EV lysates using western blotting. Syntenin-1 serves as the loading control. (d) Analysis of vesicle surface APEX2 activity. Biotinylated EVs were stained with streptavidin-AF647 conjugates and analyzed using single-vesicle flow cytometry. (e) Percentage of double position (DP, Biotin^+^mNG^+^) vesicles in total engineered EVs (mNG^+^). (f) Mean fluorescence unpaired t test. ns: non-significant; *: p < 0.05; ***: p < 0.001.

Human kidney embryonic (HEK-293T) cells were transduced with lentiviral vectors encoding the six constructs above and selected using puromycin to establish stable cell lines. At first, we examined the expression levels and activity of APEX2 at the cellular level before collecting EVs for downstream characterization. Cells were biotinylated after adding biotin-phenol and hydrogen peroxide and then stained with AF647-labeled streptavidin conjugates (Supplementary Figure 1a). Flow cytometry analysis showed that almost all cells were double positive for mNG and surface biotin, indicating effective construct expression and APEX2 activity (Supplementary Figure 1b).

Next, we harvested small EVs from the stable cell lines using tangential flow (TFF) and spin-filtration. We confirmed that the EV preparations had a median diameter of 120 nm (Supplementary Figure 2a) and were negative for the exclusion marker calnexin (Supplementary Figure 2b). Subsequently, we examined APEX2 activity at the vesicular level (Figure 2b). In the first method, EVs were biotinylated and the whole lysate was analyzed for the presence of biotin via western blot analyses. Interestingly, TSPAN2-APEX2-mNG (T2AM) exhibited the highest biotin expression, followed by CD63-APEX2-mNG (CAM), APEX2-PTTPIP-mNG (APM), and APEX2-LAMP2-mNG (ALM); however, APEX2-FPRP-mNG (AFM) had biotinylation levels comparable to that of the negative control (CD63-mNG, CM; Figure 2c). Since biotin-phenol and hydrogen peroxide can penetrate the vesicle membrane, biotinylation observed through western blotting might be driven by intravesicular APEX2. To specifically examine the activity of surface APEX2, we stained vesicle surface biotin with streptavidin-AF647 conjugates and detected intact EVs using nano-flow cytometry (Figure 2b). All constructs, except CM and AFM, lead to detectable levels of engineered vesicles (mNG-positive; Figure 2d). Moreover, T2AM and CAM EVs had the highest levels of surface biotinylation, with approximately 20% of mNG-positive vesicles being biotin-positive (Figure 2e) and an approximately 3.5-fold increase in mean AF647 fluorescence intensity over the negative control (Figure 2f). Thus, CD63 and TSPAN2 were selected as the top two scaffold proteins for surface display of APEX2 on EVs in subsequent analyses.

### Vesicle surface APEX2 labels EV surface proteins

To analyze the EV integral surfaceome, EVs were produced from CAM and T2AM stable cells cultured in serum-free Opti-MEM media and biotinylated in phosphate buffers containing biotin-phenol and H2O2 (Figure 3a). Following the removal of unreacted biotin-phenol using size exclusion chromatography columns, EVs were lysed to release biotinylated proteins. Then, streptavidin beads were applied to pull down biotinylated proteins for MS analysis using label-free quantification (LFQ) techniques. A technical negative control was included where the biotin-phenol in the biotinylation step was omitted.

**Figure 3.**
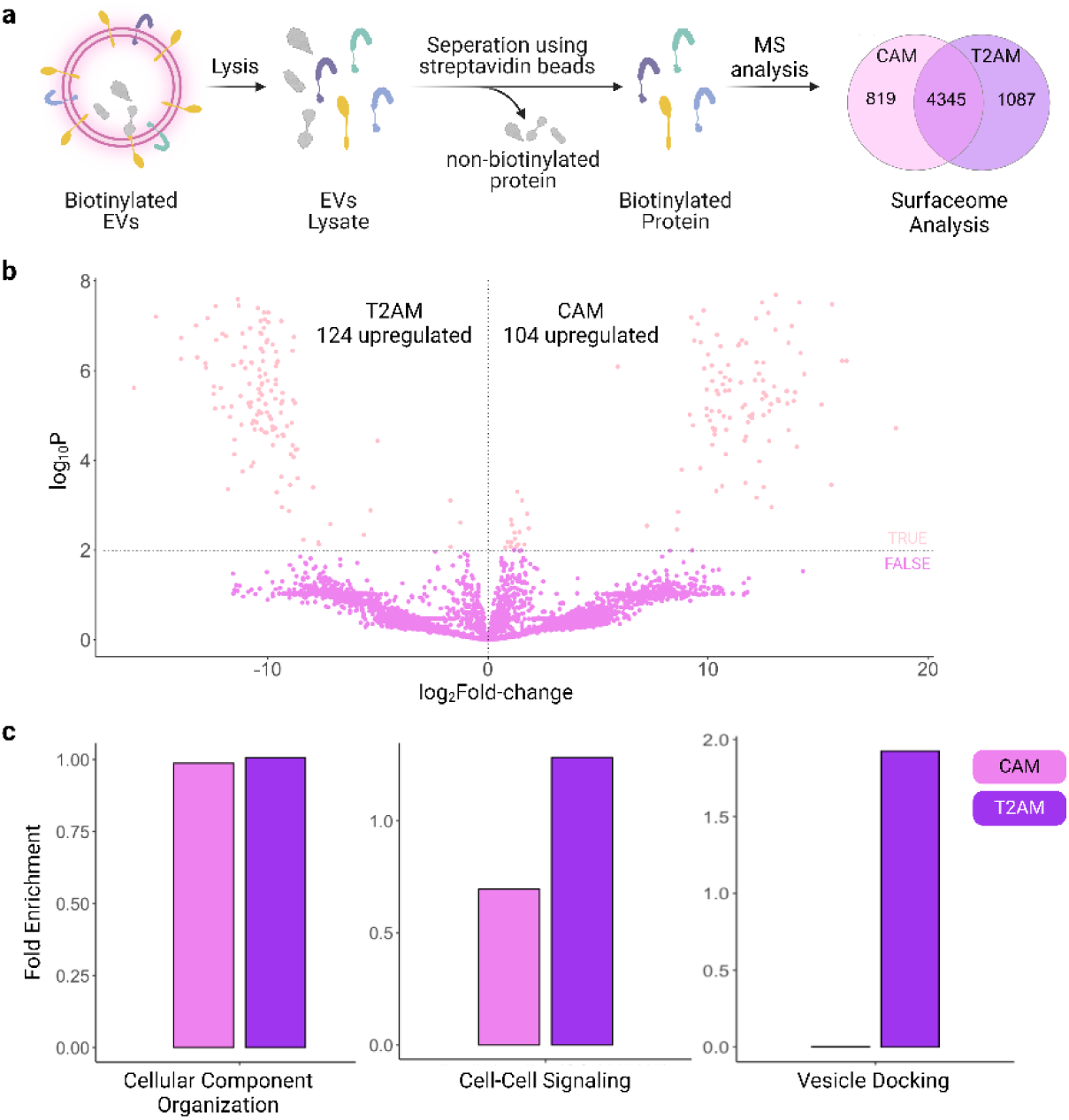
Analysis of EV integral surface proteome. (a) Nascent EVs were biotinylated by adding biotin-phenol and H2O2 and lysed to release cargo proteins. Biotinylated proteins were enriched using streptavidin beads and analyzed using conventional mass spectrometry. Venn diagram of biotinylated proteins in CAM and T2AM EVs is shown. (b) Volcano plot of biotinylated proteins in CAM and T2AM EVs. P ≤ 0.01 was considered significant. (c) Gene ontology term fold enrichment for proteins specifically enriched for each of the constructs analyzed.

Thousands of proteins, including Syntenin-1, which is primarily located inside vesicles ^38^, were identified in the negative controls (Supplementary Figure 3a), which suggested non-specific protein binding to streptavidin beads. We filtered out non-specific proteins by applying background correction and defined biotinylated proteins as those with a higher peak area than those of the negative controls. Following filtering, a total of 5,164 and 5,432 human proteins were included in the surfaceome datasets for nascent CAM and T2AM EVs, respectively. Between the two datasets, 4,345 proteins were shared, including two reported EV surface proteins, GAPDH^39^ and CD81^40^ (Figure 3a).

Next, we applied differential analysis of CAM and T2AM EVs surfaceome to gain insight into subpopulation-specific proteins. Aligning with our previous observations^37^, TSPAN2 had low basal expression levels in CAM EVs while TSPAN2-engineered EVs minimally expressed CD63 (Supplementary Figure 3b). Of the 6,023 surface proteins, we found 104 and 124 proteins were significantly (P ≤ 0.01) upregulated in CAM and T2AM EVs, respectively (Figure 3b). Gene ontology (GO) analysis of differentially expressed proteins indicated that CAM-enriched surface proteins were mostly involved in the nitrogen compound, primary, and macromolecule metabolic processes (Supplementary Figure 3c), whereas T2AM-enriched ones were involved in biogenesis and the positive regulation of cell growth (Supplementary Figure 3d). In addition, we found that T2AM surface proteins showed a higher propensity for cell-cell signaling and vesicle docking than CAM surface proteins (Figure 3c). At the protein level, integrin alpha-1 (coding gene ITGA1), a receptor for laminin and collagen, was significantly higher in T2AM EVs (Supplementary Figure 3e). Overall, the surface display of APEX2 using CD63 and TSPAN2 as scaffolds resulted in the successful labeling of the surfaceome of nascent EVs.

### EV integral surfaceome contains novel scaffolds for bioengineering

The surface display of proteins-of-interest (POI) is useful for producing EVs with decoy receptors and targeting molecules. The extracellular loop of tetraspanin proteins can tolerate the insertion of POI with proximal termini (such as antibody Fc region binder^41^, albumin binder^42^, eGFP^36^, APEX2), but not with distal termini. An alternative strategy for surface display is the fusion of POI to the extracellular termini of single-pass transmembrane proteins, such as Lamp2b^43^, FPRP^32^, PTTG1^34^, TNFR^33^, PDGFR^44^or their variants, but the efficacy of this method is highly variable as shown in this and other studies. Thus, the identification of next-generation scaffold proteins for surface display would be highly instructive.

We hypothesized that the APEX2-labelled surface proteome could potentially be exploited as a source of EV-sorting scaffold proteins. A few selection criteria were set up to shortlist candidate proteins: (1) a molecular weight less than 75 kDa to facilitate protein expression; (2) at least one transmembrane domain to exclude peripherally associated proteins; and (3) at least one extracellular terminal for genetic fusion (Figure 4a). Based on these, we identified six proteins for further investigation, including COPT1 (high affinity copper uptake protein 1), MFAP3 (microfibril-associated glycoprotein 3), PAQR1 (adiponectin receptor protein 1), PLVAP (plasmalemma vesicle-associated protein), STX4 (syntaxin-4), and S29A1 (equilibrative nucleoside transporter 1). Interestingly, the six candidates were not among the most abundant surface proteins according to the proteomics results (Figure 4b) and have molecular weights ranging from 21 kDa to 50 kDa.

**Figure 4.**
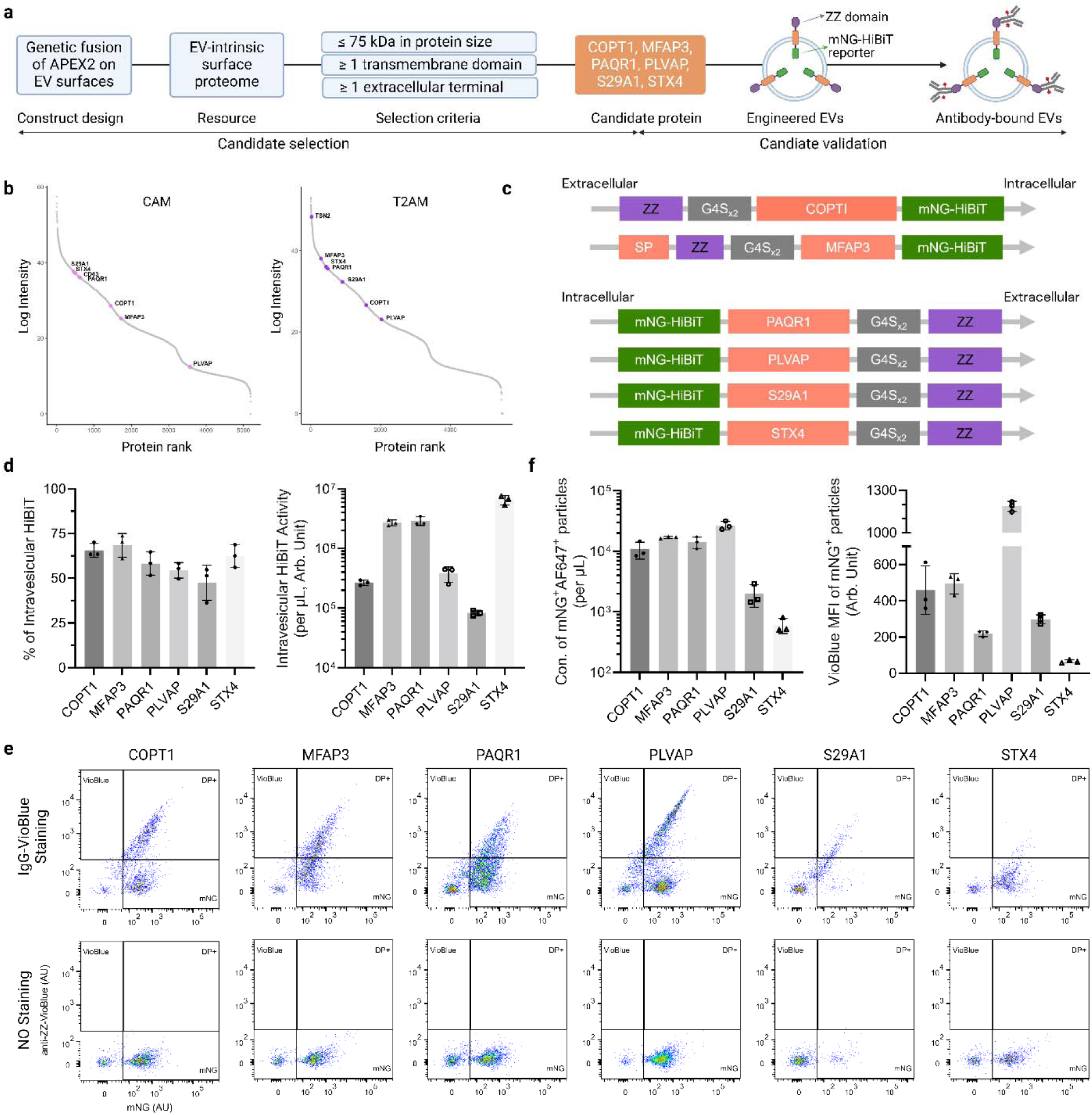
Identification of novel transmembrane domains for EV bioengineering. (a) Workflow to select candidate scaffolds from EV surface proteins and validate them at the vesicle level. (b) Protein abundance rank plot according to EV surfaceome. (c) Construct design to validate scaffold candidates. ZZ domain was fused to the candidate through a G4S linker. (d) Quantification of HiBiT activity inside engineered vesicles. (e) EVs were stained with isotype antibody-VioBlue conjugates and analyzed using single-vesicle flow cytometry. (f) Quantification of VioBlue^+^mNG^+^ vesicles. Mean ± standard deviation.

To test their potential for surface display, we cloned the ZZ domain that binds antibody Fc regions^41^ to the extracellular terminal (downstream of the signal peptide if present) and the mNG-HiBiT hybrid reporter at the other terminal for quantification (Figure 4c). HEK-293T cells were transfected with plasmids encoding the designed constructs to produce engineered EVs. Transfection efficiency was around 30% at the cellular level, with approximately 5-20% of cells being double positive for the ZZ domain and mNG (Supplementary Figure 4). The HiBiT tag enabled our quantification of the total amount of intravesicular vs. secreted fusion proteins. Approximately 50% (range 47.5-68.4%) of HiBiT signals were intravesicular for all constructs, while STX4 had the highest levels of intravesicular HiBiT signals (Figure 4d). Next, we stained EVs with VioBlue-conjugated isotype antibodies to assess surface display efficiency (Figure 4e). All constructs produced acceptable concentrations of double positive EVs (6.1×10^2^-2.6×10^4^ particles/µL) with PLVAP giving the best efficiency, followed by MFAP3 (Figure 4f). Interestingly, PLVAP-engineered EVs had the highest mean VioBlue fluorescence intensity, suggesting a superior copy number per vesicle. Thus, the EV surfaceome revealed by APEX2 proved a valuable source of novel scaffold proteins for EV engineering.

### Vesicle surface APEX2 labels EV corona proteins

The EV corona may play important biological roles and carry critical information for biomarker identification, underscoring the significance of the high-fidelity resolution of corona proteins. Here, we used vesicle surface APEX2 to label EV corona proteins. Nascent CAM and T2AM EVs were exposed to plasma from healthy mice in vitro before biotinylation (Figure 5a). Vesicle surface biotinylation was confirmed using nano-flow cytometry after staining with AF647-streptavidin conjugates (Figure 5b), but the detected labeling in plasma was not as strong as in phosphate buffers (Supplementary 5a). Since biotin-phenol and H_2_O_2_ are diffusible, we inferred that the decrease in labeling strength was the most likely due to the poor diffusion of AF647-streptavidin conjugates (around 60 kDa) within the EV corona, rather than lower biotinylation efficacy.

**Figure 5.**
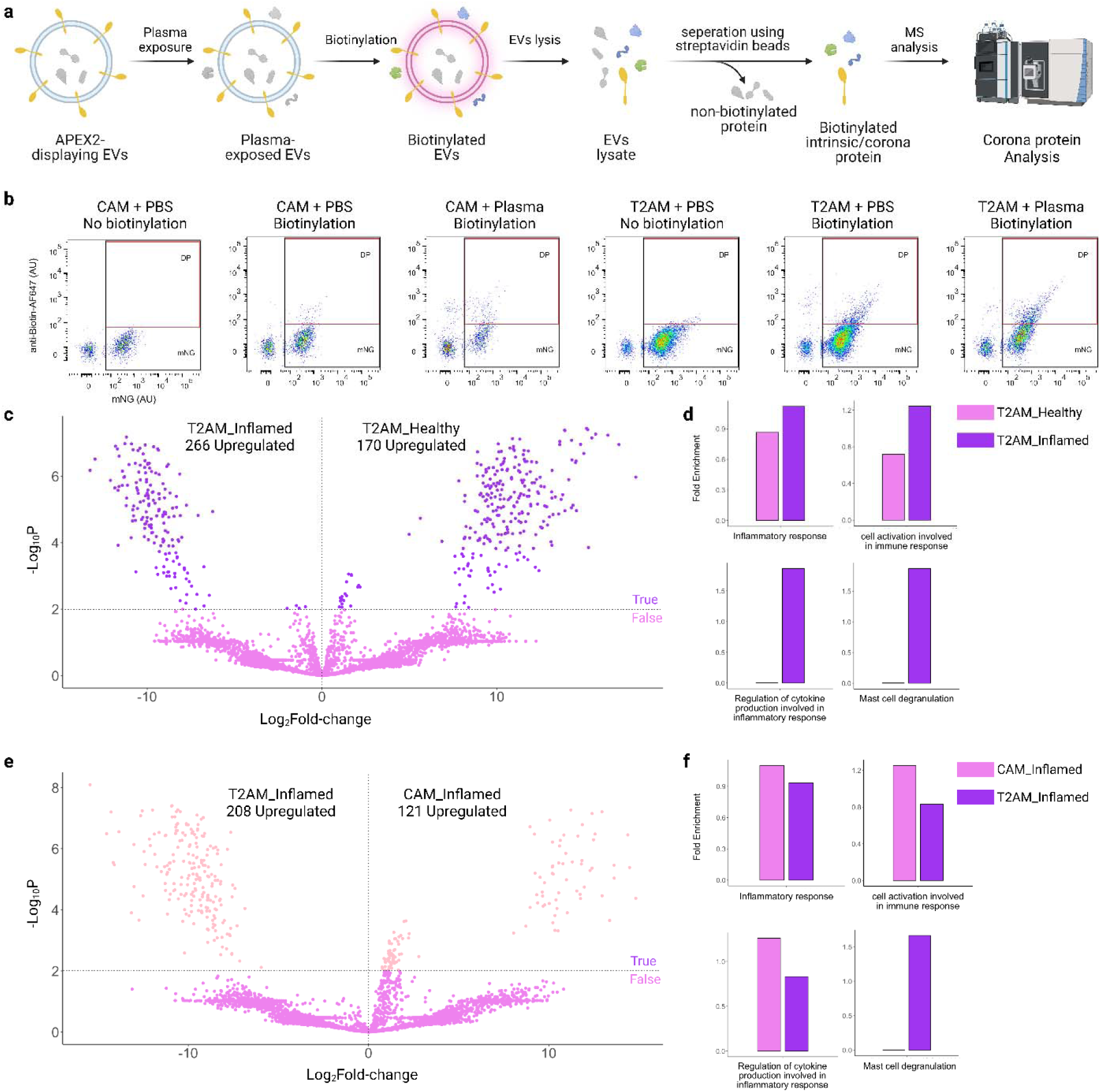
Proteomic analysis of 232 EV corona after exposure to mouse plasma. (a) Nascent EVs were mixed with mouse plasma and biotinylated. The mixture was lysed to release cargo proteins and separated with streptavidin beads to enrich biotinylated proteins for MS analysis. (b) Analysis of vesicle surface APEX2 activity in nascent and plasma-exposed EVs (c, e) Volcano plot of biotinylated proteins in CAM (c) and T2AM (e) EVs. FC, fold-change. P ≤ 0.01 was considered significant. (d, f) GO enrichment analysis of differentially expressed proteins in CAM (d) and T2AM (f) EVs.

Corona formation is contingent on matrix type. To reveal how disease context affects corona components, we compared the surfaceome of T2AM EVs following incubation with serum from healthy mice (T2AM-Healthy) or mice with lipopolysaccharide-induced systemic inflammation (T2AM-Inflamed) (Figure 5a). As noted earlier, non-biotinylated negative controls were included to account for non-specific binding to streptavidin beads. Mass spectrometry analysis identified 4,960 and 5,231 mouse proteins in the corona datasets for T2AM-Healthy and T2AM-Inflamed, respectively (Supplementary 5b). A majority (4,112) of proteins are shared between the conditions, including abundant plasma proteins such as lipoprotein and Complement factors. Differential abundance analysis of the plasma-exposed T2AM EV corona indicated that 170 and 266 proteins were significantly (P ≤ 0.01) down-regulated and up-regulated in T2AM-Inflamed samples relative to T2AM-Healthy, respectively (Figure 5c). Among the top up-regulated corona proteins in T2AM-Inflamed samples were Gapdh (glyceraldehyde-3-phosphate dehydrogenase), Mpp1 (membrane-associated guanylate kinase P55 scaffold protein 1), ribosomal proteins, Sphk1 (sphingosine Kinase 1), and Urod (uroporphyrinogen decarboxylase), while the top down-regulated protein was Srprb (signal recognition particle receptor subunit beta) (Supplementary Figure 5c). Moreover, GO analysis of the differentially expressed proteins revealed that T2AM-Inflamed samples were more enriched with proteins involved in the inflammatory response, cell activation involved in immune response, regulation of cytokine production, and mast cell degranulation (Figure 5d) than their healthy counterparts.

In addition, we compared EV subpopulation-specific corona proteins by incubating CAM or T2AM EVs with plasma from inflammatory mouse models. A total of 4,669 proteins were discovered in the CAM-Inflamed surfaceome dataset, with 3,968 proteins shared with T2AM-Inflamed samples (Supplementary Figure 5b). A total of 329 proteins were significantly differentially expressed between CAM-Inflamed and T2AM-Inflamed samples (Figure 5e). Among the top up-regulated corona proteins in CAM-Inflamed samples were Ap1m1 (adaptor related protein complex 1 subunit mu 1) and Usp5 (ubiquitin specific peptidase 5), while Ca1 (carbonic anhydrase 1), Cmpk2 (cytidine/uridine monophosphate kinase 2), Itih4 (inter-alpha-trypsin inhibitor heavy chain 4), Lamp2, Mri1 (methylthioribose-1-phosphate isomerase 1), Napa (NSF attachment protein alpha), and Rnf11 (ring finger protein 11) were most down-regulated (Supplementary Figure 5d). GO analysis of the differentially expressed proteins points to the enrichment of cell activation in immune response, regulation of cytokine production, and inflammatory response, but the depletion of mast cell degranulation components in CAM-Inflamed samples (Figure 5f).

Lastly, we compared the overall effect of the EV subpopulation or plasma type on corona formation using D-means clustering analysis. Interestingly, plasma types were found to make a higher contribution than molecular subpopulation (384.2 vs. 342.8) to corona formation (Supplementary Figure 5e). Overall, the surface display of APEX2 using CD63 and TSPAN2 as scaffolds robustly labels EV corona proteins after plasma exposure.

### Vesicle surface APEX2 preferentially labels target cells

Identifying EV target cells is fundamental for elucidating EVs’ mechanisms of action and developing therapeutic EVs. Currently, this is mainly achieved through labeling EVs with reporter proteins (e.g., fluorescent and bioluminescent proteins, and nucleases) and/or synthetic tracers and quantifying cellular uptake^26^, which innately neglects target cells that respond in a contact-dependent manner. Here, we determined if vesicle surface APEX2 could label cell surface interactors during the limited time of contact between EVs and cells. Consequently, we designed a proof-of-concept experiment to test if T2AM EVs label target cells over non-target cells (Figure 6a).

**Figure 6.**
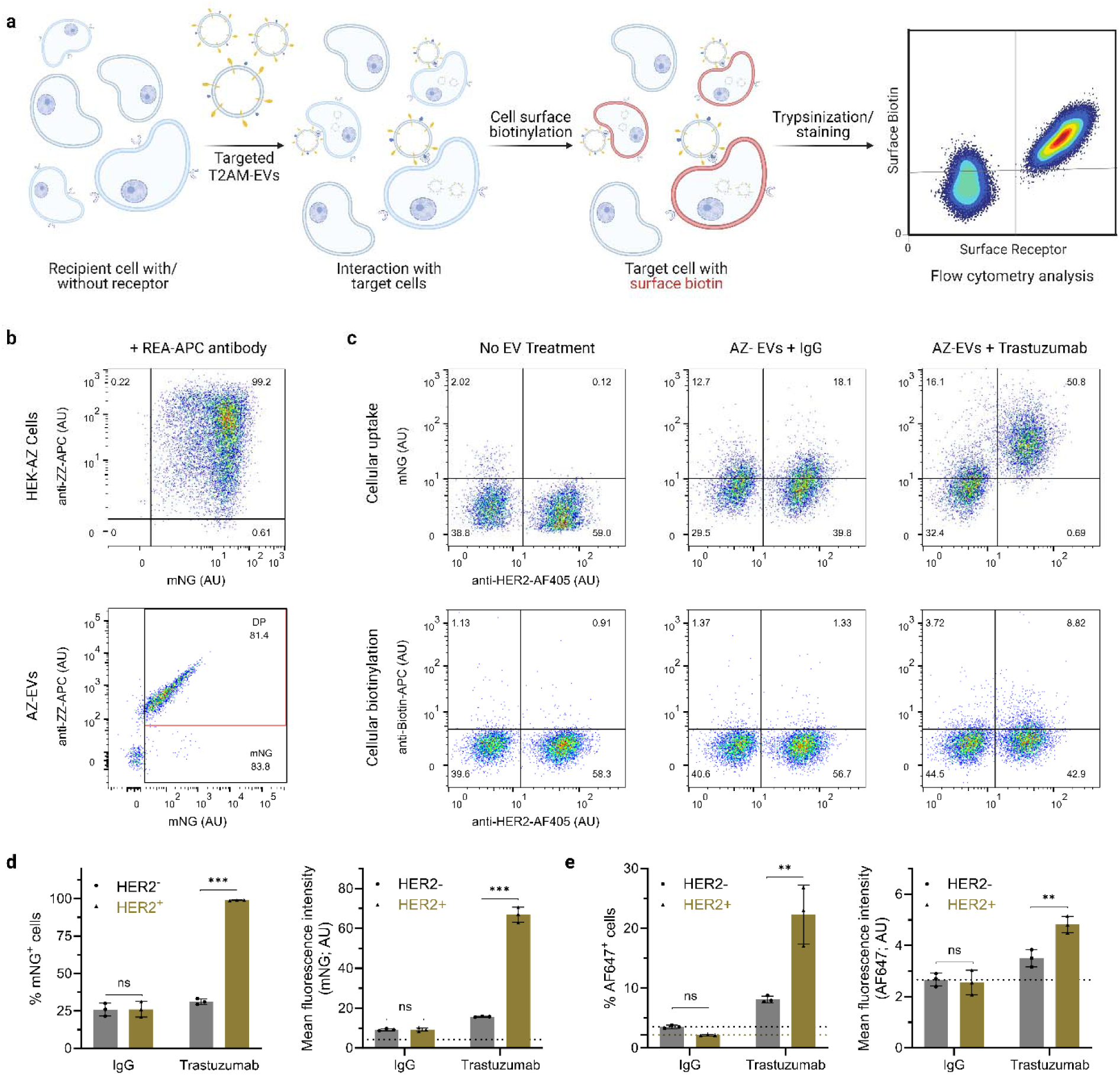
Surface 307 display of APEX2 on HER2-targeted EVs. (a) HEK-293T cells were transduced to co-express TSPAN2-ZZ-Nluc and TSPAN2-APEX2-mNG (AZ cells) and the generated AZ EVs were incubated with trastuzumab to construct HER2-targeted EVs. Recipient cells were a mixture of B16F10 cells with or without HER2. After EV treatment, cells were biotinylated, trypsinized, and quantified for cellular mNG and surface biotin levels. (b) Dot-plot of AZ cells and EVs after staining with APC-conjugated isotype antibodies. (c) Dot-plot of B16F10 cells after EVs treatment, biotinylation, and staining with streptavidin-AF647 conjugates. (d) Quantification of cellular uptake of EVs in target (HER2^+^) and non-target (HER2^-^ cells. (e) Quantification of cell surface biotinylation of EVs in target (HER2^+^) and non-target (HER2^-^) cells. Mean ± standard deviation. Two-tailed parametric unpaired t test. ns: non-significant; **: p < 0.01; ***: p < 0.001.

Our group has recently developed HER2 (human epidermal growth factor receptor 2)-targeted EVs through surface display of the ZZ domain that binds anti-HER2 trastuzumab.^41^Similar to the original design, the ZZ domain was cloned at the large extracellular loop of TSPAN2 (T2Z). To produce HER2-targeted T2AM EVs, HEK-T2AM cells were additionally transduced to stably express T2Z, and the EVs thereof (named AZ) were coated with trastuzumab. Flow cytometry analysis revealed that T2AM and T2Z fusion proteins are highly co-localized at the cellular and single-vesicle levels (Figure 6b). Subsequently, we established the recipient cell model by co-culturing (1:1) mouse melanoma B16F10 cells with or without HER2 expression. Recipient cells were treated with trastuzumab or isotype-coated EVs for 2 h and biotinylated as described earlier. The cellular uptake of EVs was measured by tracking EV-derived mNG signals (Figure 6c). As expected, coating with trastuzumab, but not isotype antibody significantly increased EV uptake by HER2-positive cells relative to HER2-negative cells (Figure 6d). More importantly, coating with trastuzumab resulted in a significant increase in cell surface biotin levels in HER2-positive cells (Figure 6c), with a 2.75-fold higher number of stained cells and a 1.38-fold higher intensity than in HER2-negative cells (Figure 6e). Taken together, these results demonstrate that the vesicle surface APEX2 can be leveraged to identify target cells.

## DISCUSSION

In this study, we have developed an effective proximity labeling platform for the high-fidelity characterization of EV integral and corona proteins and the mapping of EV-interacting cells. Moreover, we report six novel scaffold proteins amenable for simultaneous surface display and intravesicular loading of cargo proteins. Classic scaffold proteins that are widely used (e.g., CD9, CD63, CD81^45^, Lamp2^43^, and ARRDC1^46^) were derived from our prior biological understanding of EV biogenesis, while novel scaffolds were identified via proteomic data mining of whole EV lysate (for PRGFRN^32^, BASP1^32^, and PTTG1IP^34^) or the screening of tetraspanin proteins (for TSPAN2 and TSPAN3^37^). Our study provides a novel workflow to identify scaffold proteins for genetic engineering of EVs.^47^ This workflow builds on EV surfaceome data rather than that from whole EV lysate and is thus better suited for screening EV scaffold proteins for surface display. This study relies on the surfaceome of CD63 and TSPAN2-engineered EVs from HEK-293T cells. In the future, it would be valuable to delve into the surfaceome of alternative EV subpopulations and producer cells, to potentially identify additional scaffold proteins. Notably, the low abundance of EVs in liquid biopsies poses a significant hurdle in developing diagnostic biomarkers^48^, calling for a specific and efficient enrichment method. Apart from their use in biomedical engineering, the newly identified EV integral surface proteins might serve as biomarkers and antigens for the immunoaffinity-based capture of EVs.

Corona proteins associate with the EV surface via various mechanisms and can therefore be specifically isolated in several ways. These include the use of digestive enzymes (trypsin, proteinase K, and RNase)^49,50^, cation chelators (ethylenediaminetetraacetic acid) to disrupt electrostatic interactions^51^, and reducing agents (tris(2-carboxyethyl)phosphine) to cleave disulfide bonds^52^. As such, proteins susceptible to these treatments are often characterized as surface proteins. Alternative studies have characterized the EV surfaceome using tethering PL enzymes or applying membrane-impermeable chemical probes (like NHS-biotin^28^ to label protein amines and aminooxy-biotin^53^ to label glycoproteins). A common procedure of these methods is the removal of confounders (such as free proteins and non-vesicular particles) to yield a relatively pure batch of EVs. However, the shear force introduced by this purification step might irreversibly deplete corona proteins, particularly loosely associated ones. In this study, plasma-exposed EVs were not purified, but instead immediately biotinylated prior to molecular characterization, thereby providing a snapshot of the near-native state of the corona. Owing to the use of gravity-based SEC columns to remove unreacted biotin-phenol, the possibility of biotinylated proteins dissociating from the EV surface during this procedure was presumed to be much less due to the absence of shear centrifugal force. Using our protocol as a reference, one could potentially compare the relative efficiency of common isolation methods, such as precipitation, ultracentrifugation, and ultrafiltration, in preserving the molecular signatures of EVs in biological fluids.

EVs affect recipient cells by means of intracellular delivery of vesicular cargo molecules as well as interfacial interactions without internalization. Notably, the efficiency of intracellular delivery is modulated by these surface interactions between EVs and cells.^54^ Here, we found that vesicle surface APEX2 preferentially labels EV target cells over non-target cells in vitro. Complementary to available tools that determine the cellular uptake of EVs^26^, this study presents a novel technique for detecting the repertoire of EV-interacting cells irrespective of cellular uptake. EV integral and corona proteins and target cell-derived surface molecules are essential mediators of cellular tropism, and further studies are warranted to annotate these molecules. Generally, the results of our study provide a novel framework for the annotation of protein interactions between EVs and recipient cells. This insight will be essential to understanding EV biology and to exploit them as biomarkers or drug delivery vehicles.

## METHODS

### Plasmids

Codon-optimized DNA sequences coding for proteins-of-interest were ordered commercially (Integrated DNA Technologies and Twist Biosciences) and cloned downstream of the CAG promoter into the pLEX vector using restriction cloning strategies. Lentiviral transfer plasmids were synthesized by cutting transgenes from pLEX expression plasmids and inserting them into p2CL9IPwo5 backbone (plasmid was kindly given by H. Hanenberg, University Hospital Essen, Germany). All expression cassettes were confirmed by Sanger sequencing (Eurofins Genomics). Plasmids are available from the corresponding author upon request.

### Cell culture

HEK-293T (ATCC, CRL-3216) and B16F10 (ATCC, CRL-6475) cells were maintained in high glucose DMEM (Gibco, 41966-029) media supplemented with 10% fetal bovine serum (FBS; Gibco, 10270-106) and 1% anti-anti (Gibco, 15240). All cells were cultured in humidified incubators with 37°C and 5% CO_2_.

### Plasmid transfection

HEK-293T were seeded in 15-cm Petri dishes containing 20 mL media and transfected upon confluency reaching approximately 60%. 30 µg plasmid was complexed with 45 µg polyethyleneimine (Polysciences; 24765-1) in 4 mL Opti-MEM (Gibco, 31987-047) and added to each dish. Media was changed to Opti-MEM 6 h later and cells were cultured for an additional 48 h before harvesting conditioned media.

### Lentivirus production and transduction

HEK-293T cells were seeded in T-175 flasks containing 20 mL DMEM and transfected upon confluency reaching approximately 60%. Cells were co-transfected overnight with 22 µg transfer plasmids, 22 µg helper plasmids pCD/NL-BH, and 3.5 µg envelope plasmids pcoPE (encoding the human foamy virus envelope protein) that were pre-complexed with 135 µg polyethyleneimine in 4 ml Opti-MEM. The media was changed to full DMEM supplemented with 10 mM sodium butyrate (Sigma-Aldrich) for 8 h to induce gene expression and changed again to DMEM. The conditioned media was collected 22 h later and filtered through 0.45 µm filters. The filtrate was further spun at 25,000 g for 90 min at 4 ℃ and the pellet was suspended in 1 ml Iscove’s modified Dulbecco’s media supplemented with 20% FBS and 1% antibiotic antimycotic. The viruses were kept at -80 ℃ until use.

HEK-293T cells were seeded in 6-well plates until being approximately 60% confluent and then transduced with lentiviral particles overnight. To make stable cell pools, cells were passed a minimum of five times under antibiotics selection (4 µg/ml puromycin or 10 µg/ml blasticidin).

### Extracellular vesicles production

HEK-293T-derived stable cell lines were cultured in 15-cm Petri dishes containing 20 mL DMEM until confluence reached approximately 90%. Media was changed to Opti-MEM and cells were cultured for an additional 48 h. The conditioned media was sequentially centrifuged (700 g for 5 min and 2000 g for 10 min) and filtered (0.2 µm) to deplete cells, cellular debris, and large particles. Then, filtrate was diafiltrated and concentrated to roughly 80 mL using the KrosFlo KR2i TFF System (Repligen, US) with 300 kDa cut-off hollow fiber filters (Spectrum Labs, D06-E300-05-N) at a flow rate of 70 mL/min (transmembrane pressure at 3.0 psi and shear rate at 3700 sec^-1^). EVs were further concentrated till approximately 0.5 mL using an Amicon Ultra-15 spin-filter with 10 kDa molecular weight cut-off (Millipore, UFC901024) and stored at -80 ℃ in PBS-HAT buffer until use.

### Particle analysis

EVs were diluted in 200 nm-filtered PBS if required and analyzed using ZetaView Twin (Particle Metrix) at the following settings: sensitivity of 75, shutter of 130, 11 position measurement scans per sample, 2 cycles. Particle size and concentration were derived using the default setting.

### Cellular and vesicular biotinylation

HEK and B16F10 cells were cultured in 24-well plates till confluency reached around 60%. B16F10 cells were treated with 1e10 EVs for 2 h and rinsed with PBS twice. To trigger cellular biotinylation, 5 µl biotin-phenol (50 mM in DMSO, Sigma-Aldrich, SML2135-50MG) was added per well and left for 30 min. Then, 5 µl H_2_O_2_ (100 mM) was added and left for 1 min at room temperature. Excess H_2_O_2_ was quenched using 0.5 ml antioxidant mixture (50 mM Trolox, 100 mM sodium ascorbate, 100 mM sodium azide).

Vesicular biotinylation was performed similarly. Briefly, EVs (5e10 particles in 200 µl) were incubated with a 200 µL matrix (PBS or plasma) in a tube at 37 ℃ for 30 min. Four µl biotin-phenol (50 mM in DMSO) was added and incubated at 37 ℃ for another 30 min. Then, 4 µl H_2_O_2_(100 mM in PBS) was added and left at room temperature (RT, 22 ℃) for 5 min. Excess H_2_O_2_ was quenched using a 12 µL antioxidant mixture as before. Afterwards, EVs were stained with AF647-labeled streptavidin conjugates before flow cytometry analysis.

### Size exclusion chromatography

EVs were purified using gravity-based size exclusion chromatography (SEC) columns (Izon, SP1) according to the manufacturer’s instructions. The first 3 ml eluate was discarded as the void volume, while the next 2.5 ml eluate that was enriched with EVs was collected. The EV fraction was further concentrated using 10 kDa cut-off spin filters and stored at -80 ℃.

### Mouse plasma

All mouse experiments were performed following the ethical permission granted by Swedish Jordbruksverket (permit No. 20281-2021). Balb/c mice (8-week-old) were bought from Charles River and housed in an animal facility for at least one week before use according to standard routines (ambient temperature: 20-22 ℃ °C, humidity: 45-55%, dark/light cycle: 12/12 h). Mice were intraperitoneally injected with lipid polysaccharide at 4 mg/kg. Blood was collected 5 h later into heparinized tubes and centrifuged at 2000g for 10 min. The supernatant was collected and stored at -80 ℃ until use.

### Western blotting

EVs (2×10^9^ particles in 15 µL) were mixed with a 5 µL sample buffer (NP0007, Invitrogen) and heated at 70 ℃ for 10 min. The lysate was loaded onto a NuPAGE Novex 4–12% Bis-Tris Protein Gel (Invitrogen, NP0335BOX) and separated in NuPAGE MES SDS running buffer (Invitrogen, NP0002) at 120 V for 2 h. Proteins were transferred to a nitrocellulose membrane (Invitrogen, IB23001) using the iBlot system. The membrane was immersed in blocking buffer (LI-COR, 927-60004) for 1 h at room temperature and incubated with primary antibodies (1:1000 dilution for anti-Calnexin [ThermoFisher, PA5-19169], anti-Syntenin-1 [Origene, TA504796], Streptavidin-680RD [926-68079, LI-COR]) overnight at 4 ℃. Afterwards, the membrane was rinsed using TBS supplemented with 0.1% Tween 20 (TBS-T) for 3 times and stained with corresponding secondary antibodies (925-68070, 926-32350, LI-COR; 1:10,000 dilution) for 1 h at RT. Eventually, the membrane was rinsed with TBS-T 3 times, once with PBS, and visualized on the Odyssey infrared imaging system (LI-COR, US).

### Streptavidin pulldown

Biotinylated EVs were added to a lysis buffer containing 0.05% lauryl maltose neopentyl glycol (LMNG), 0.1% Triton-X, and a protease inhibitor cocktail (Roche). Lauryl maltose neopentyl glycol (LMNG) was added to prevent protein adsorption to the tube.^55^ The samples were further sonicated using Bioruptor for 3 min with a 15-sec run and 15-sec intervals. To pull down biotinylated proteins, the EV lysate was incubated with 20 µl of streptavidin magnetic beads (Pierce, 88817) at 4 °C for 2 h under rotation. Protein-bound beads were sequentially rinsed with RIPA buffer (once), 1 M KCl solution (once), 0.1M Na_2_CO_3_solution (once), and 50 mM NH₄HCO₃ buffer (twice). For every rinse step, the beads were well suspended in a 1 ml buffer and rotated for 1 min before separating on a magnetic stand.

To liberate proteins for proteomics analysis, the beads were suspended in 200 µl 50 mM NH₄HCO₃ buffer supplemented with 0.02% LMNG. Then, proteins were reduced and alkylated with 10 mM tris(2-carboxyethyl)phosphine and 50 mM 1M chloroacetamide at 37 °C for 30min. Proteins were digested using 0.5 µg trypsin (Promega) at 37 °C overnight under shaking (>1000 RPM). The next day, the supernatant was transferred to new tubes and acidified with 20 µl 10% trifluoroacetic acid. Peptide samples were desalted using the SDB-RPS StageTip.^56^

### LC/MS/MS analysis

LC/MS/MS analysis was performed on an Orbitrap Eclipse mass spectrometer (Thermo Fisher Scientific) equipped with a FAIMSpro interface, combined with a Vanquish Neo UHPLC pump (Thermo Fisher Scientific). The mobile phases consisted of (A) 0.1% formic acid and (B) 0.1% formic acid and 80% acetonitrile (ACN). Peptides were loaded on a self-made 18 cm fused-silica emitter (100 µm inner diameter) packed with ReproSil-Pur C18-AQ (1.9 µm, Dr. Maisch, Ammerbuch, Germany), and separated by a linear gradient for 60 min (5–40% B over 45 min, 40–99% B over 5 min, and 99% B for 10 min) at the flow rate of 250 nl/min. FAIMS compensation voltages (CVs) were fixed to –45. MS scanning was performed in the data-independent acquisition (DIA) mode using the Orbitrap analyzer. MS1 scans were performed in the range of 350 to 1000 m/z (resolution = 120,000, maximum injection time = 45 ms, and automatic gain control = 300%). In the following MS/MS scans, the precursor range was set to 500 to 740 m/z, and 20 scans were acquired with the isolation window of 12 m/z, with HCD normalized collision energy of 27 (resolution = 50,000, injection time = 86 ms, auto gain control = 1000%, first mass = 120 m/z).

Raw files were processed with default settings using DIA-NN (v.1.8.1)^57^ to perform a library-free search against the UniProt/SwissProt human and/or mouse database (downloaded from UniProt February 2024) combined with the contaminant database obtained from MaxQuant.^58^ The following additional options were used: --protein-qvalue 0.01; --peak-translation; --mass-acc-cal 10; --relaxed-prot-inf; --matrix-qvalue 0.01; --matrix-spec-q; and --top 4.

Subsequently, LFQ intensities from biotinylated samples were background subtracted from non-biotinylated samples for further analyses. The data was then log-transformed and normalized by median centering. The missing values were replaced by a normal distribution with a downshift of 1.8 SDs and a width of 0.3 SDs. To identify proteins upregulated in each of the specified samples, a linear model was fit through the data, and the empirical Bayes method was used to moderate the standard errors of the estimated log-fold changes using the limma package.^59^ A false discovery rate (FDR) of p_adjusted_ < 0.01 was used here to minimize the risk of Type 1 error (i.e. false positives). This method results in more inference and improved power for small datasets.

Gene ontology (GO) enrichment analyses for proteins upregulated for each construct were analyzed using the Panther Classification system.^60^ The data was analyzed using Fischer’s Exact test and the GO overrepresentation annotation set. A control gene set was constructed from the consolidation of all upregulated and the fold enrichments displayed for each term were based on the division of the FE of the group of interest by that of the control group. All analyses were conducted using R (v 4.2.3) and all plots were created using ggplot2.

### Flow cytometry for cells

Cell surface biotinylated proteins were stained with 100 µl streptavidin-AF647 conjugates (10 ng/ml) at room temperature for 30 min. After trypsinization, cells were resuspended in 100 µL of PBS containing 2% FBS. DAPI (4,6-diamidino-2-phenylindole) was added to all samples to exclude dead cells. The samples were measured with a MACSQuant Analyzer 16 cytometer (Miltenyi, Germany). Data was analyzed with FlowJo software (version 10.6.2) and doublets were excluded by forward scatter area versus height gating (gating strategy available in Supplementary Figure 6).

### Flow cytometry for single vesicle

25 µL EVs (maximum concentration 1 × 10^10^ particles/mL) were stained with 5 µl streptavidin-AF647 conjugates (6 ng/ml) or VioBlue-isotype antibodies (6 ng/ml) at room temperature overnight and diluted by 2,00-fold in PBS-HAT buffer. Data was acquired on the Amnis CellStream instrument (Luminex) equipped with 405, 488, 561, and 642 nm lasers. Samples were measured with FSC turned off, SSC laser set to 40%, and all other lasers set to 100%. EVs were defined as SSC (low) by using mNG-tagged EVs as biological reference material, and regions to quantify mNG^+^ or AF647^+^ fluorescent events were set according to unstained non-fluorescent samples and single fluorescence positive mNG-tagged reference EV controls. Data was analyzed using FlowJo software (version 10.6.2, gating strategy available in Supplementary Figure 6).

### Luciferase detection assay

To detect HiBiT in conditioned media, 25 μl samples were added into white-walled 96-well plates along with 25 µl PBS or Triton X-100 solution. The plate was shaken horizontally for 5 min. 50 µL of ready-to-use HiBiT Lytic Detection mixture (Promega; N3040) was added to each well. After incubation at room temperature under horizontal shaking for 10 min, the plate was immediately measured.

### Statistics & Reproducibility

Results were shown as mean (± standard deviation) of biological replicates. The two-sided Student’s t test was used to compare the mean difference between two groups. No statistical method was used to predetermine the sample size. No data were excluded from the analyses. The Investigators were not blinded to allocation during experiments and outcome assessment.

## Supplementary Information

Supplementary data are provided in this paper.

## Acknowledgments

SEA is supported by the European Research Council (ERC) under the European Union’s Horizon 2020 research and innovation program (DELIVER, grant agreement No. 101001374; EXPERT, grant agreement No. 825828), Swedish Foundation of Strategic Research (FormulaEx, SM19-0007), Cancer Foundation (grant No. 21-1762-Pj-01-H), Swedish Research Council (grant No. 4–258/2021). KI is supported by the Japan Science and Technology (JST) PRESTO (JPMJPR18H2).

## Author Contribution

WZ conceptualized the study, performed experiments, and wrote the manuscript. MM performed bioinformatic analysis. WH performed experiments. KI designed and performed the proteomic analysis. SEA and DWH supervised the project and provided funding. All authors edited the manuscript before submission.

## REFERENCE

1. Couch, Y. et al. A brief history of nearly EV-erything - The rise and rise of extracellular vesicles. J Extracell Vesicles 10, (2021).

2. Margolis, L. & Sadovsky, Y. The biology of extracellular vesicles: The known unknowns. PLoS Biol 17, e3000363-(2019).

3. van Niel, G. et al. Challenges and directions in studying cell–cell communication by extracellular vesicles. Nat Rev Mol Cell Biol 23, 369–382 (2022).

4. Tóth, E. Á. et al. Formation of a protein corona on the surface of extracellular vesicles in blood plasma. J Extracell Vesicles 10, e12140 (2021).

5. Buzás, E. I., Tóth, E. Á., Sódar, B. W. & Szabó-Taylor, K. É. Molecular interactions at the surface of extracellular vesicles. Semin Immunopathol 40, 453–464 (2018).

6. Buzas, E. I. Opportunities and challenges in studying the extracellular vesicle corona. Nat Cell Biol 24, 1322–1325 (2022).

7. Liam-Or, R. et al. Cellular uptake and in vivo distribution of mesenchymal-stem-cell-derived extracellular vesicles are protein corona dependent. Nat Nanotechnol (2024) doi:10.1038/s41565-023-01585-y.

8. Seras-Franzoso, J. et al. Extracellular vesicles from recombinant cell factories improve the activity and efficacy of enzymes defective in lysosomal storage disorders. J Extracell Vesicles 10, e12058 (2021).

9. Harmati, M., Bukva, M., Böröczky, T., Buzás, K. & Gyukity-Sebestyén, E. The role of the metabolite cargo of extracellular vesicles in tumor progression. Cancer Metastasis Rev 40, 1203–1221 (2021).

10. Pham, T. T. et al. Endocytosis of red blood cell extracellular vesicles by macrophages leads to cytoplasmic heme release and prevents foam cell formation in atherosclerosis. J Extracell Vesicles 12, 12354 (2023).

11. Valadi, H. et al. Exosome-mediated transfer of mRNAs and microRNAs is a novel mechanism of genetic exchange between cells. Nat Cell Biol 9, 654–659 (2007).

12. Crewe, C. et al. Extracellular vesicle-based interorgan transport of mitochondria from energetically stressed adipocytes. Cell Metab 33, 1853–1868.e11 (2021).

13. Hansen, A. S. et al. T-cell derived extracellular vesicles prime macrophages for improved STING based cancer immunotherapy. J Extracell Vesicles 12, 12350 (2023).

14. Roefs, M. T. et al. Cardiac progenitor cell-derived extracellular vesicles promote angiogenesis through both associated- and co-isolated proteins. Commun Biol 6, 800 (2023).

15. Chen, G. et al. Exosomal PD-L1 contributes to immunosuppression and is associated with anti-PD-1 response. Nature 560, 382–386 (2018).

16. Poggio, M. et al. Suppression of Exosomal PD-L1 Induces Systemic Anti-tumor Immunity and Memory. Cell 177, 414–427.e13 (2019).

17. Zhong, W. et al. Tumor-Derived Small Extracellular Vesicles Inhibit the Efficacy of CAR T Cells against Solid Tumors. Cancer Res 83, 2790–2806 (2023).

18. Rausch, L. et al. Phosphatidylserine-positive extracellular vesicles boost effector CD8+ T cell responses during viral infection. Proceedings of the National Academy of Sciences 120, e2210047120 (2023).

19. Raposo, G. et al. B lymphocytes secrete antigen-presenting vesicles. Journal of Experimental Medicine 183, 1161–1172 (1996).

20. Blandin, A. et al. Extracellular vesicles are carriers of adiponectin with insulin-sensitizing and anti-inflammatory properties. Cell Rep 42, 112866 (2023).

21. Jin, S. et al. Astroglial exosome HepaCAM signaling and ApoE antagonization coordinates early postnatal cortical pyramidal neuronal axon growth and dendritic spine formation. bioRxiv 2023.02.14.528554 (2023) doi:10.1101/2023.02.14.528554.

22. Ter-Ovanesyan, D. et al. Improved isolation of extracellular vesicles by removal of both free proteins and lipoproteins. Elife 12, e86394 (2023).

23. Karimi, N. et al. Detailed analysis of the plasma extracellular vesicle proteome after separation from lipoproteins. Cellular and Molecular Life Sciences 75, 2873–2886 (2018).

24. Wang, T. & Turko, I. V. Proteomic Toolbox To Standardize the Separation of Extracellular Vesicles and Lipoprotein Particles. J Proteome Res 17, 3104–3113 (2018).

25. Heidarzadeh, M., Zarebkohan, A., Rahbarghazi, R. & Sokullu, E. Protein corona and exosomes: new challenges and prospects. Cell Communication and Signaling 21, 64 (2023).

26. Boudna, M. et al. Strategies for labelling of exogenous and endogenous extracellular vesicles and their application for in vitro and in vivo functional studies. Cell Communication and Signaling 22, 171 (2024).

27. Qin, W., Cho, K. F., Cavanagh, P. E. & Ting, A. Y. Deciphering molecular interactions by proximity labeling. Nat Methods 18, 133–143 (2021).

28. Kirkemo, L. L. et al. Cell-surface tethered promiscuous biotinylators enable comparative small-scale surface proteomic analysis of human extracellular vesicles and cells. Elife 11, 2021.09.22.461393 (2022).

29. Li, Y., Kanao, E., Yamano, T., Ishihama, Y. & Imami, K. TurboID-EV: Proteomic Mapping of Recipient Cellular Proteins Proximal to Small Extracellular Vesicles. Anal Chem 95, 14159–14164 (2023).

30. Zheng, W. et al. Surface display of functional moieties on extracellular vesicles using lipid anchors. Journal of Controlled Release 357, 630–640 (2023).

31. Naito, M., Nagashima, K., Mashima, T. & Tsuruo, T. Phosphatidylserine Externalization Is a Downstream Event of Interleukin-1β–Converting Enzyme Family Protease Activation During Apoptosis. Blood 89, 2060–2066 (1997).

32. Dooley, K. et al. A versatile platform for generating engineered extracellular vesicles with defined therapeutic properties. Molecular Therapy 29, 1729–1743 (2021).

33. Gupta, D. et al. Amelioration of systemic inflammation via the display of two different decoy protein receptors on extracellular vesicles. Nat Biomed Eng 5, 1084–1098 (2021).

34. Martin Perez, C., et al. An extracellular vesicle delivery platform based on the PTTG1IP protein. bioRxiv 2023–2028 (2023).

35. Hung, V. et al. Spatially resolved proteomic mapping in living cells with the engineered peroxidase APEX2. Nat Protoc 11, 456–475 (2016).

36. Corso, G. et al. Systematic characterization of extracellular vesicles sorting domains and quantification at the single molecule–single vesicle level by fluorescence correlation spectroscopy and single particle imaging. J Extracell Vesicles 8, 1663043 (2019).

37. Zheng, W. et al. Identification of scaffold proteins for improved endogenous engineering of extracellular vesicles. Nat Commun 14, 4734 (2023).

38. Kugeratski, F. G. et al. Quantitative proteomics identifies the core proteome of exosomes with syntenin-1 as the highest abundant protein and a putative universal biomarker. Nat Cell Biol 23, 631–641 (2021).

39. Dar, G. H. et al. GAPDH controls extracellular vesicle biogenesis and enhances the therapeutic potential of EV mediated siRNA delivery to the brain. Nat Commun 12, 6666 (2021).

40. Mathieu, M. et al. Specificities of exosome versus small ectosome secretion revealed by live intracellular tracking of CD63 and CD9. Nat Commun 12, 4389 (2021).

41. Wiklander, O. P. B. et al. Antibody-displaying extracellular vesicles for targeted cancer therapy. Nat Biomed Eng (2024) doi:10.1038/s41551-024-01214-6.

42. Liang, X. et al. Extracellular vesicles engineered to bind albumin demonstrate extended circulation time and lymph node accumulation in mouse models. J Extracell Vesicles 11, e12248 (2022).

43. Alvarez-Erviti, L. et al. Delivery of siRNA to the mouse brain by systemic injection of targeted exosomes. Nat Biotechnol 29, 341–345 (2011).

44. Ohno, S. et al. Systemically Injected Exosomes Targeted to EGFR Deliver Antitumor MicroRNA to Breast Cancer Cells. Molecular Therapy 21, 185–191 (2013).

45. Andreu, Z. & Yáñez-Mó, M. Tetraspanins in Extracellular Vesicle Formation and Function. Front Immunol 5, (2014).

46. Wang, Q. et al. ARMMs as a versatile platform for intracellular delivery of macromolecules. Nat Commun 9, 960 (2018).

47. Richter, M., Vader, P. & Fuhrmann, G. Approaches to surface engineering of extracellular vesicles. Adv Drug Deliv Rev 173, 416–426 (2021).

48. Liang, Y., Lehrich, B. M., Zheng, S. & Lu, M. Emerging methods in biomarker identification for extracellular vesicle-based liquid biopsy. J Extracell Vesicles 10, e12090 (2021).

49. Choi, D. et al. Quantitative proteomic analysis of trypsin-treated extracellular vesicles to identify the real-vesicular proteins. J Extracell Vesicles 9, 1757209 (2020).

50. Xu, R. et al. Surfaceome of Exosomes Secreted from the Colorectal Cancer Cell Line SW480: Peripheral and Integral Membrane Proteins Analyzed by Proteolysis and TX114. Proteomics 19, 1700453 (2019).

51. Rai, A., Fang, H., Claridge, B., Simpson, R. J. & Greening, D. W. Proteomic dissection of large extracellular vesicle surfaceome unravels interactive surface platform. J Extracell Vesicles 10, e12164 (2021).

52. Santucci, L. et al. Biological surface properties in extracellular vesicles and their effect on cargo proteins. Sci Rep 9, 13048 (2019).

53. Chauhan, S. et al. Surface Glycoproteins of Exosomes Shed by Myeloid-Derived Suppressor Cells Contribute to Function. J Proteome Res 16, 238–246 (2017).

54. Hallal, S., Tűzesi, Á., Grau, G. E., Buckland, M. E. & Alexander, K. L. Understanding the extracellular vesicle surface for clinical molecular biology. J Extracell Vesicles 11, (2022).

55. Konno, R. et al. Universal Pretreatment Development for Low-input Proteomics Using Lauryl Maltose Neopentyl Glycol. Molecular & Cellular Proteomics 23, (2024).

56. Kulak, N. A., Pichler, G., Paron, I., Nagaraj, N. & Mann, M. Minimal, encapsulated proteomic-sample processing applied to copy-number estimation in eukaryotic cells. Nat Methods 11, 319–324 (2014).

57. Demichev, V., Messner, C. B., Vernardis, S. I., Lilley, K. S. & Ralser, M. DIA-NN: neural networks and interference correction enable deep proteome coverage in high throughput. Nat Methods 17, 41–44 (2020).

58. Tyanova, S., Temu, T. & Cox, J. The MaxQuant computational platform for mass spectrometry-based shotgun proteomics. Nat Protoc 11, 2301–2319 (2016).

59. Ritchie, M. E. et al. limma powers differential expression analyses for RNA-sequencing and microarray studies. Nucleic Acids Res 43, e47–e47 (2015).

60. Mi, H. et al. Protocol Update for large-scale genome and gene function analysis with the PANTHER classification system (v.14.0). Nat Protoc 14, 703–721 (2019).

