## Supplementary Figure for "Surface display of proximity labeling enzymes on extracellular vesicles for surfaceome and target cell mapping"

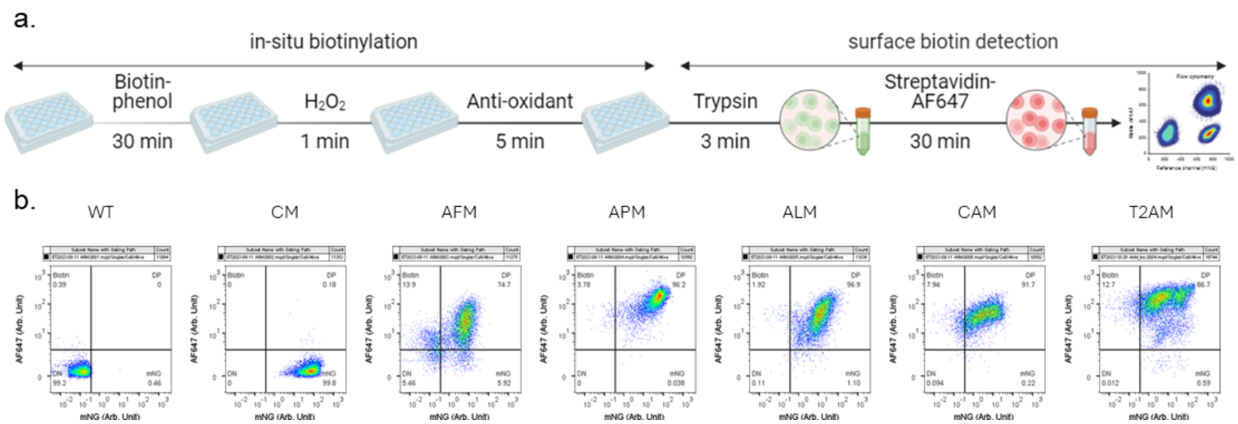


**Supplementary Figure 1. Validation of APEX2 activity on cell surface.** (a) Cells were grown in multi-well plates and treated with biotin-phenol for 30 min. Biotinylation was triggered via a short treatment with $H_2O_2$ and quenched with antioxidants. Afterwards, cells were trypsinized and stained with streptavidin-AF647 conjugates before flow cytometry analysis. Dead cells were gated out using DAPI. (b) Quantification of APEX2 construct expression and activity in cells. The parent gate was single live cells.

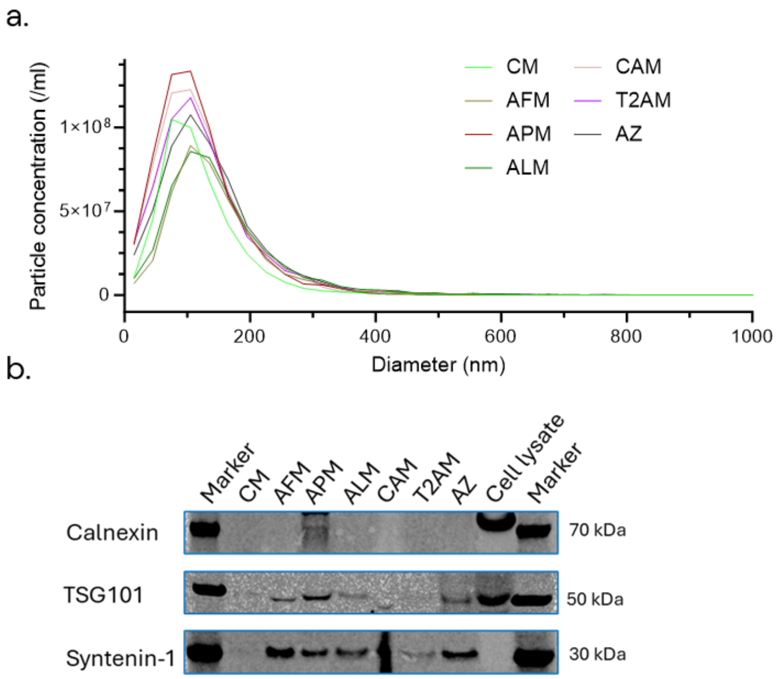


**Supplementary Figure 2. Physiochemical characterization of EVs.** (a) Size distribution analysis using ZetaView. (b) Blots against common vesicle markers and exclusion markers.

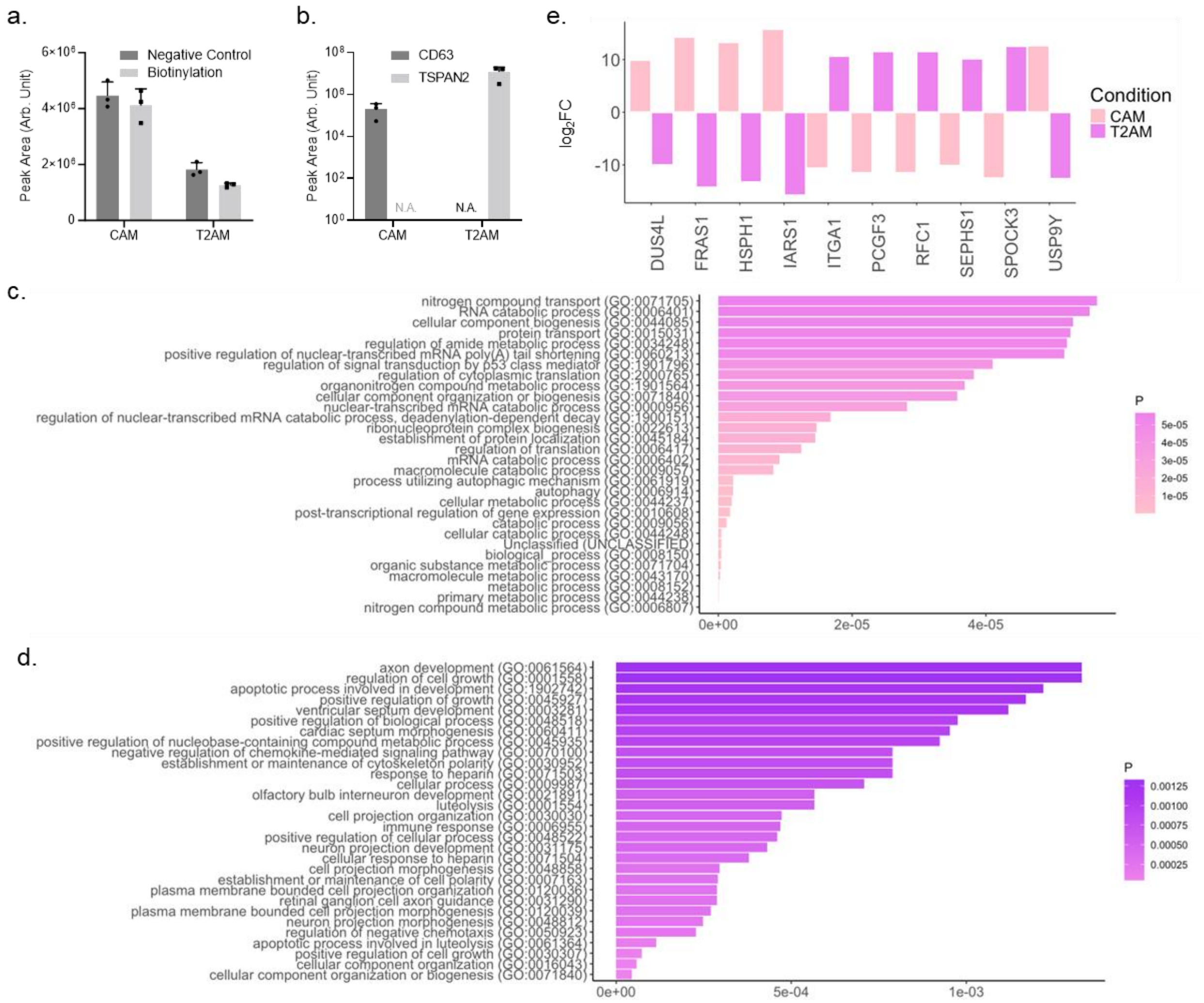


**Supplementary Figure 3. Proteomic analysis of EV-intrinsic surface proteins.** (a) Syntenin-1 protein peak area in CAM and T2AM EVs. (b) CD63 and TSPAN2 protein peak area in CAM and T2AM EVs. (c) Gene ontology analysis of CAM EVs surfaceome. (d) Gene ontology analysis of T2AM EVs surfaceome. (e) Relative abundance ($log_2$fold-change) of top ten differentially expressed proteins.

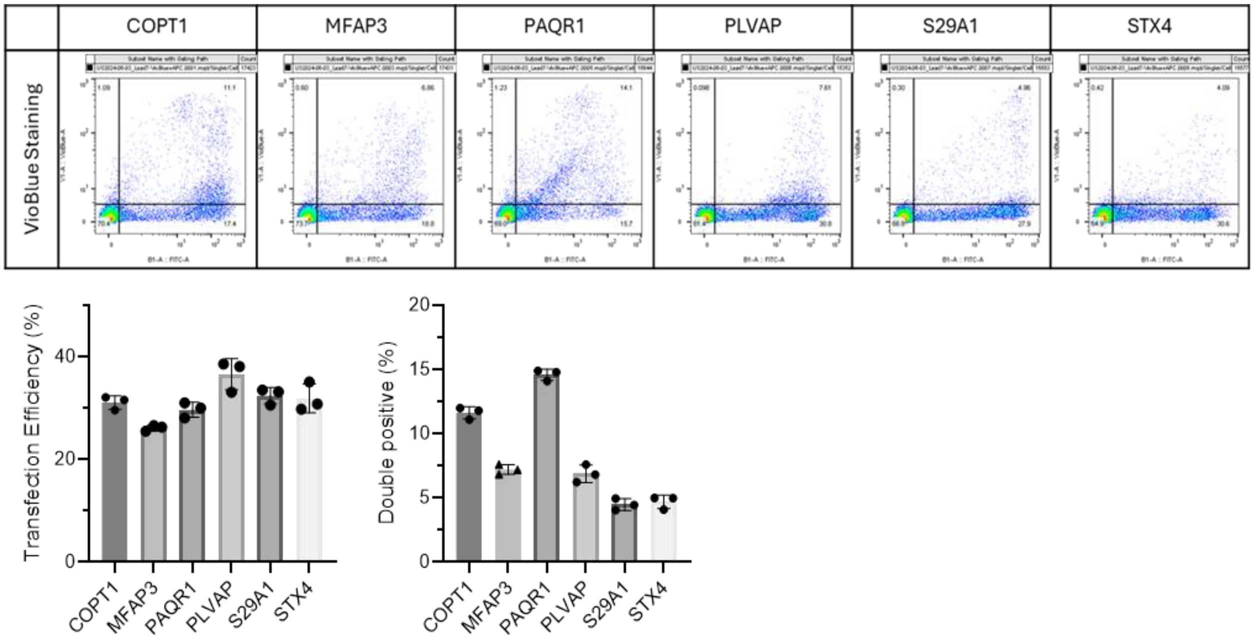


**Supplementary Figure 4. Surface display construct expression on transfected cells.** HEK-293T cells were transfected with the constructs and stained with isotype antibody-VioBlue conjugates. Transfection efficiency was the percentage of $mNG^+$ cells. The parent gate was single live cells. Mean ± standard deviation.

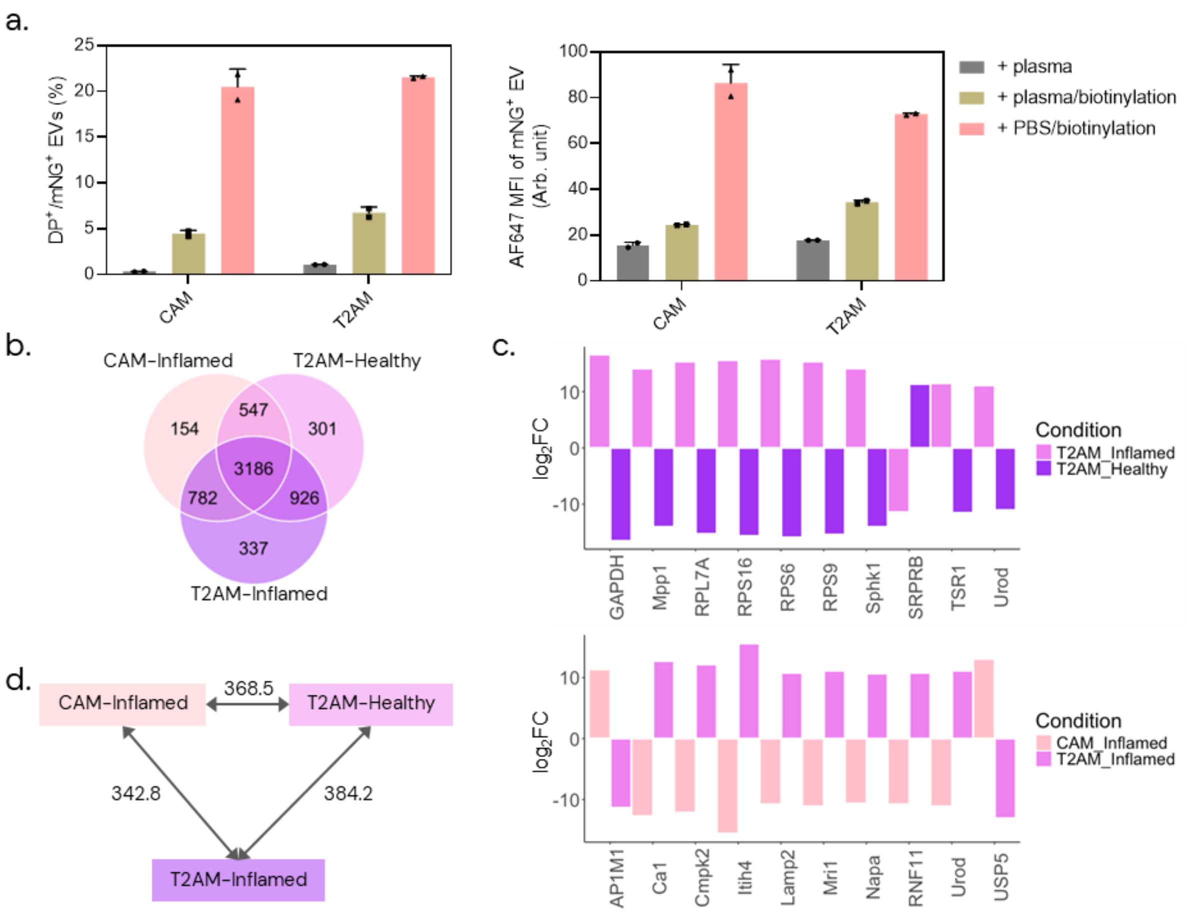


**Supplementary Figure 5. Proteomic analysis of EV corona proteins after exposure to mouse plasma.** (a) Analysis of vesicle surface APEX2 activity in nascent and plasma-exposed EVs. (b) Venn diagram of biotinylated mouse proteins in plasma-exposed CAM and T2AM EVs. (c) Relative abundance ($log_2$fold-change) of top ten differentially associated mouse proteins. (d) Distances between cluster centroids obtained from K-means clustering analysis.

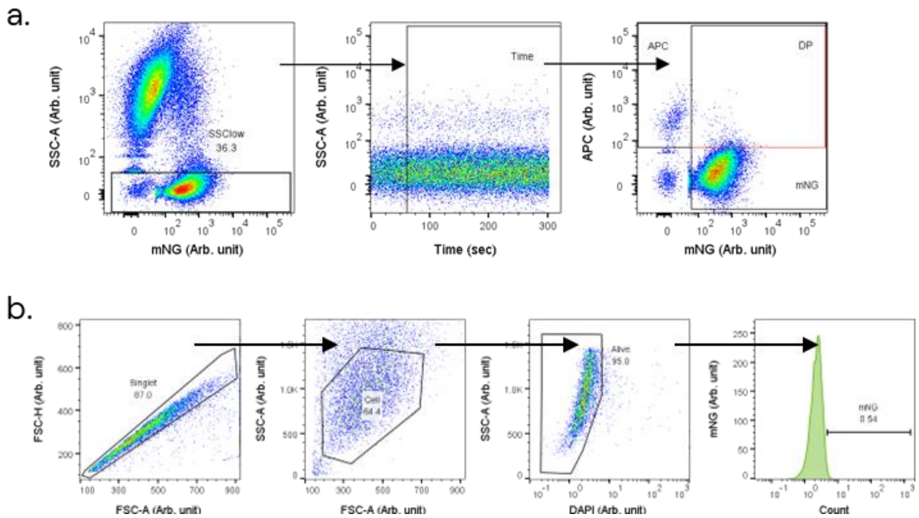


**Supplementary Figure 6. Gating strategy for EVs (a) and cells (b) in flow cytometry.**
